# LC-MS Based Draft Map of the *Arabidopsis thaliana* Nuclear Proteome and Protein Import in Pattern Triggered Immunity

**DOI:** 10.1101/2021.02.16.431396

**Authors:** Mohamed Ayash, Mohammad Abukhalaf, Domenika Thieme, Carsten Proksch, Mareike Heilmann, Martin Schattat, Wolfgang Hoehenwarter

**Author notes:** Address correspondence to: Wolfgang Hoehenwarter, Leibniz Institute of Plant Biochemistry, Biochemistry of Plant Interactions Department, Proteome Biology of Plant Interactions Research Group. Equal contribution (Mohamed Ayash, Mohammad Abukhalaf).

## Abstract

Despite its central role as the ark of genetic information and gene expression the plant nucleus is surprisingly understudied. We isolated nuclei from the *Arabidopsis thaliana* dark grown cell culture left untreated and treated with flg22 and nlp20, two elicitors of pattern triggered immunity (PTI) in plants, respectively. An LC-MS based discovery proteomics approach was used to measure the nuclear proteome fractions. An enrichment score based on the relative abundance of cytoplasmic, mitochondrial and golgi markers in the nuclear protein fraction allowed us to curate the nuclear proteome producing high quality catalogs of around 3,000 nuclear proteins under untreated and both PTI conditions. The measurements also covered low abundant proteins including more than 100 transcription factors and transcriptional co-activators. Protein import into the nucleus in plant immunity is known. Here we sought to gain a broader impression of this phenomenon employing our proteomics data and found 157 and 73 proteins to be putatively imported into the nucleus upon stimulus with flg22 and nlp20, respectively. Furthermore, the abundance of 93 proteins changed significantly in the nucleus following elicitation of immunity. These results suggest promiscuous ribosome assembly and retrograde signaling from the mitochondrion to the nucleus including Prohibitins and Cytochrome C, in the two forms of PTI.

## Introduction

Subcellular compartmentalization is a defining characteristic of eukaryotic organisms and higher cell life. Cellular organelles are membrane enclosed spaces with specific architecture and physiological milieus. They are dynamic in nature undergoing morphological changes throughout the cell cycle and in response to environmental stimuli as well as moving throughout the cytoplasm and making and breaking contact with one another for instance by way of stromules. Extensive molecular traffic between the organelles facilitates optimal biochemical metabolism and cellular function much as on an assembly line in a factory.

The plant nucleus is a very large organelle that encloses the genome, extensively reviewed by (Meier et al., 2017). The genome is scaffolded on histone proteins, the mass of which is approximately equal to the mass of DNA. The cell cycle leads to radical changes in chromatin structure and chromosome positioning and indeed cell division leads to the complete breakdown of the nuclear envelope for the mitotic spindle to access the chromosomes. These processes are all orchestrated by nuclear proteins. The plant nuclear lamina is also composed of proteins although it is not as well explored as in animals. Gene expression requires a host of DNA chromatin associated proteins, transcriptional co-activators and repressors and transcription factors. Ribosomes are assembled in nuclear sub-compartments called nucleoli. The plant nuclear envelope is populated by protein complexes that serve to control nuclear morphology, positioning and movement among other things. Thus the nucleus is populated by an estimate of several thousand proteins and the nuclear proteome in itself is highly diverse dependent on the biological cell state.

Nuclear pore complexes facilitate protein in- and export to the nucleus. In many cases proteins are the convergence points of organelle initiated second messenger or small molecule signaling, that imported into the nucleus, initiate a transcriptional response (Kmiecik et al., 2016). Both plastids and mitochondria generate reactive oxygen species (ROS) and Ca2+ fluxes that as organelle signals converge on specific kinases and transcription factors that convey the signal to the nucleus (Crawford et al., 2018). This type of retrograde signaling and sub-cellular protein trafficking between compartments has been shown to be instrumental in the cellular response to various types of abiotic stress as well as in the establishment of plant immunity.

Plant immunity is composed of multiple overlapping layers that show many of the same effects; for more information see (Zhang et al., 2020; Zhou and Zhang, 2020). The recognition of molecular patterns indicative of non-self (pathogen or microbe associated molecular patterns PAMPS or MAMPs) or of non-homeostasis self (damage associated molecular patterns DAMPs) by receptors is central to the initiation of defense. In the resistance to biotrophic pathogens, pathogens in the apoplast are first recognized by plasma membrane spanning pattern recognition receptors (PRRs) such as Leucine-rich repeat receptor like kinases (LRR-RLKs) or receptor like proteins (LRR-RPs) that recruit kinases to initiate signaling. Initial recognition and signaling is often potentiated by a host of other molecular recognition events that together propagate the signal. The best studied example of these processes is signaling downstream of the LRR-RLK FLAGELLIN SENSING 2 (FLS2) that recognizes bacterial flagellin (Gómez-Gómez and Boller, 2000). Its 22 amino acid N-terminal epitope, flg22, is a commonly used elicitor that we have also used here. The LRR-RP RLP23 is another closely related PRR that recognizes phytotoxic virulence factors ethylene-inducing peptide 1 (Nep1)-like proteins (NLPs) and initiates a similar response. Signaling is activated by perception of the characteristic 20 amino acids long peptide nlp20 (Albert et al., 2015) that we have also used in this study.

Early events following PAMP perception are the production of reactive oxygen species and Ca2+ influx. Ca2+ is essential because it directly controls many immune regulatory proteins such as calcium dependent protein kinases (CPKs) and others (Boudsocq et al., 2010). Mitogen associated protein kinase (MAPK) signaling is another central pillar of immune signaling that leads to a broad range of events by way of their phosphorylated activated substrates (Bigeard et al., 2015). Many of these are transcription factors that orchestrate reprogramming of gene expression. Phytohormones, particularly salicylic (SA), jasmonic acid (JA) and ethylene (ET) also play important roles in regulating immunity (Pieterse et al., 2012). These events are collectively termed pattern triggered immunity (PTI) and shift the plant away from homeostasis to a state of immunity that is hallmarked by the production of pathogenesis-related (PR) proteins and antimicrobial secondary metabolites.

## Methods

### Preparation of the protoplasts

Thirty mL of 5 days old *Arabidopsis thaliana* cultured cells grown in the dark was centrifuged at 805 g for 5 minutes at room temperature (RT). The pellets were resuspended in 30 mL of 0.24 M CaCl2. Then 15 mL of this suspension, 20 mL of 0.24 M CaCl2 and 15 mL of the enzyme solution (0.2 % macerozyme, 0.67 % cellulose and 0.24 M CaCl_2_) were transferred to a petri dish. The petri dish was incubated at RT overnight shaking at 45 rpm. The content of the petri dish was centrifuged at 290 g for 5 min at RT and the pellets were resuspended in 30 mL of 0.24 M CaCl_2_. The same centrifugation step was repeated but, the pellets were resuspended in 14 mL of B5 sucrose solution (0.32 % gamborg B5 medium, 1 mg/L 2, 4-D and 0.28 M sucrose at pH 5.5). The final suspension was centrifuged at 130 g for 5 minutes at RT and was left for 5-10 minutes on the rack. The floating protoplasts were collected from the top layer. Protoplast samples were supplemented with flg22 and nlp20 respectively to a concentration of 1 µM in solution and incubated for 16 hours at 18 °C. Control samples were untreated and incubated similarly. These experiments were performed three times independently (three biological replicates for each condition).

## Preparation of nuclear and cellular fractions

4 mL of protoplasts were mixed with 9 mL of NIBA (25 % v/v NIB 4x, 1mM DTT and 1 % protease inhibitor) in a falcon tube and kept on ice for 10 minutes. Triton X-100 was added to an in solution concentration of 0.1% and the suspension was gently mixed for 5 minutes. Three consecutive centrifugation steps were done each at 1000 g for 15 minutes at 4°C. After the first 2 steps the pellets were resuspended in 4 mL NIBA containing 0.1 % triton X-100. The supernatants were retained as the cellular fraction. Then the pellets were resuspended in 4 mL NIBA (washing step) and centrifuged as before. After the third step, the pellets were resuspended in 300 µL extraction buffer and transferred to a 1.5 mL Eppendorf tube (nuclear fraction, NF). NIBA 4x and extraction buffer were taken from a nuclear isolation kit (CELLYTPN1-1KT for plants, SIGMA).

### Staining and microscopy

A fluorescence microscope (Axioplan2 imaging, Carl Zeiss) with a DAPI filter was used to visualize DAPI stained nuclei. 100 µL of 5 µg/mL DAPI were added to 10 µL of the nuclear fraction and kept in darkness for 15 min. Then, 10 µL of this solution were used for microscopy.

### Extraction of nuclear proteins

200 µL of extraction buffer (containing 1% protease inhibitor cocktail, Sigma P9599) were added to the NF. The sample was vortexed at 1800 rpm for 30 min at RT and then it was sonicated in an ultrasonicator for 10 min. Then, the sample was centrifuged in a fixed rotor angle centrifuge at 12000 g for 10 min at RT. The supernatant was collected representing Nuclear Proteins.

### Extraction of cellular proteins

5 mL of CF was mixed with 45 mL of 100 mM ammonium acetate in methanol. The mixture was kept at −20°C overnight and then three centrifugation steps were done with a swinging bucket rotor centrifuge at 3200 g for 15 min at 4°C. The pellets from the first two centrifugation steps were washed with 3 mL of 20% 50 mM ammonium bicarbonate and 80% acetone and the final pellets were left to dry at RT. The dried pellets were solubilized in urea buffer (8M urea and 50 mMTris) and constituted cellular proteins.

### Western blot analysis

Five µg of the protein extracts were separated into one gradient SDS-PAGE (20-4%, Serva). The proteins were transferred to a nitrocellulose membrane using a wet blot technique (Protran, GE Healthcare). The membranes were blocked with 3% fat-free dry milk (BioRad) in TBS (50 mM Tris-HCl pH7.5, 150 mM NaCl). The blocked membranes were incubated with polyclonal antihistone H3 antibody (Agrisera) and a secondary anti-rabbit antibody coupled to HRP. Detection was performed with SuperSignal™ West Femto Maximum Sensitivity Substrate (Thermo) and the signal was recorded with the Fusion Solo S Chemiluminescence Imaging System (VWR) using a 16-bit CCD camera.

### In-solution digestion of proteins using trypsin

The protein samples were reduced by addition of DTT solution (29.9 µg/µL). Then, the samples were kept at 22°C for 1 hour shaking at 450 rpm. Samples were alkylated by the addition of iodoacetamide solution (35.9 µg/µL) and kept at 22°C for 1 hour shaking at 450 rpm in darkness. Again, the reducing solution was added to samples and was kept at 22°C for 1 hour with shaking. 50 mM ammonium bicarbonate pH 8.5 was added to each sample. Trypsin (0.2 µg/µL) was added to a ratio of 1:50. Protein digestion was allowed to proceed overnight at 37°C shaking at 750 rpm. The next day, samples were dried in a vacuum concentrator.

### STAGE-Tip C18 peptide desalting (Stop-and-Go Extraction)

Dried peptides were dissolved in 200 µL 0.1% formic acid (FA). Desalting was done using In house C18-STAGE-Tips. STAGE-Tips were inserted into 1.5 mL Eppendorf tubes and conditioned with 100 µL 80% acetonitrile (ACN), 0.1% formic acid (FA) by centrifugation at 1500 g for 2 minutes at RT. Then, they were equilibrated two times with 100 µL 0.1% FA with subsequent centrifugation at 1500g for 2 minutes at RT. The dissolved peptides were applied to STAGE-Tips and centrifuged twice at 1500 g for 2 min at RT. The flow-throughs were discarded. The STAGE-Tips were washed twice with 100 µL 0.1 % FA and centrifuged as before. The flow-throughs were discarded. STAGE-Tips were inserted into new 1.5 mL Eppendorf tubes and elution was done twice by adding 50 µL of 80% ACN, 0.1% FA followed by centrifugation at 1500 g for 1 min at RT. The eluates (peptides) were combined and dried in a vacuum concentrator

### Liquid chromatography and mass spectrometry (LC-MS/MS)

The dried peptides were dissolved in 10 µL of 5% ACN, 0.1% FA. The samples were analyzed on a Q Exactive Plus mass spectrometer equipped with an EASY nanoLC-1000 liquid chromatography system (both from ThermoFisher Scientific). A flow rate of 250 nL/min was used. Peptides were separated using an analytical column ES803A (ThermoFisher) and a gradient increasing from 5% to 40% of solvent B (ACN in 0.1% FA) in 540 minutes followed by 13 minutes of isocratic flow at 80% of solvent B (for Cellular proteins). On the other hand, the nuclear proteins peptide samples were separated using a gradient inclining from 5% to 35% of solvent B (ACN in 0.1% FA) in 450 minutes followed by 20 minutes of incline to 80% solvent B and finally fixed at 80% solvent B for 70 minutes. The spray voltage was set to 1.9 KV and the capillary temperature to 275°C.

A Data-Dependent Acquisition (DDA) scan strategy was used, where one MS full scan was followed by up to 10 MS2 scans of product ions from the 10 most abundant precursor ions. The MS full scan parameters were: AGC target 3E+06, resolution 70, 000 and max injection time (IT) 100 ms. The MS2 parameters were: resolution 17,500, Max IT 50 ms, dynamic exclusion duration 40 s, ACG target 5E+04 and isolation window 1.6 m/z.

### Identification and quantification of peptides and proteins

Peptide and by inference protein identification was done by matching the MS raw data with *in silico* generated peptide ion *m/z* and MS2 spectral peak lists. The TAIR10 protein database (14486974 residues, 35394 sequences) was searched using the Mascot search engine V2.5.1 coupled to the Thermo proteome discoverer 2.1.1.21. The enzyme specificity was set to trypsin with tolerance of 2 missed cleavages. Ion m/z error tolerance was set to 5 ppm and 0.02 Da for precursor and fragment ions respectively. Carbamidomethylation of cysteine was set as a static modification and oxidation of methionine as a variable modification. Peptide spectral match (PSM), peptide and protein level false discovery rates (FDR) were determined by a decoy database search. A significance threshold of α= 0.01 was used for PSM and peptide level identifications. For the protein level, α of 0.05 was tolerated. The PSM count was used as protein abundance quantitative index (PQI). Protein grouping was inferred based on the principal of parsimony and only master proteins (protein group member that best explains the set of peptides used for inference) were retained. In the case of duplicate gene models producing individual master proteins, the first gene model was retained (this was the case in less than 1% of master proteins).

### Bioinformatics data analysis

Gene ontology analysis of the curated nuclear proteome was performed using the DAVID Bioinformatics resources 6.8 (Huang et al., 2009a; Huang et al., 2009b) using default parameters. *Arabidopsis Thaliana* was used as background and TAIR_ID was used as identifier. Functional annotation chart was created with threshold of count: 2 and ease:0.1. Proteins annotated to the nucleus, with GOTERM_CC_DIRECT, were further clustered using high classification stringency. Subcellular location of the curated nuclear proteome was checked with SUBA4 (Hooper et al., 2017) using experimental locations inferred by fluorescent protein or MS/MS studies (retrieval from SUBA4 was done on January 2020 and rechecked on February 2021). LOCALIZER 1.0.4 (Sperschneider et al., 2017) was used to predict organelle subcellular localization by searching for targeting sequences such as NLS in protein primary structure. To further evaluate the possible biological role of the proteins in significant functional categories, their AGI codes were used to query the STRING database (Szklarczyk et al., 2019) for physical interaction setting the stringency to highest confidence interactions, which we have shown to be true positive previously (Hoehenwarter et al., 2013), using experiments, databases, co-occurrence and co-expression as interaction sources and showing only interactions between proteins in the input set. All raw and meta data will be made available to the public via ProteomeXchange/PRIDE.

### Collective data analysis

Mean PSM value of each protein in all measurements of the nuclear fraction (µNp_n_) and the cellular fraction (µCp_n_) was calculated. These values were used to formulate two scores, the nuclear and cellular enrichment scores (Npf_n_ and Cpf_n_) that express the ratio of the abundance of protein (n) in the nuclear and the cellular fraction respectively (equation 1 and 2).

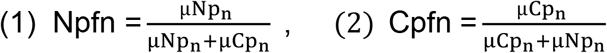

### Statistical data analysis

The matrix of curated nuclear proteins PQI (PSM) values of all samples was imported to Perseus software v.1.6.6.0 (Tyanova et al., 2016). The PQI values were grouped into 3 groups (control, flg22 and nlp20). Proteins that did not have a value in at least 5 of the 6 measurements of at least one group were discarded. The individual measurements (columns) were unit vectors normalized. Multiple sample test (ANOVA) was performed for the 3 groups in order to assess the significance of changes in abundance between conditions using permutation-based FDR multiples testing correction with an FDR significance threshold α of 0.05 and 250 permutations. Post hoc test (FDR= 0.05) was performed to identify the significant group pairs. Proteins with statistically significant changes in their abundance were kept and their values Z-score transformed. Hierarchical clustering was performed using Pearson correlation as distance measure for row clustering and Spearman correlation for columns.

## Results

In this study we set out to produce a high quality draft catalog of the *Arabidopsis thaliana* nuclear proteome based on isolation of nuclei and mass spectrometric (MS) measurement of nuclear protein fractions. Beyond this we investigated the quantitative changes in protein abundance in the nuclear proteome in the three biological scenarios (control, flg22 and nlp20), and we took first steps towards gaining better understanding of protein import into the nucleus and retrograde signaling in two related forms of PTI elicited using flg22 and nlp20 respectively. To do this we chose to expose *Arabidopsis* protoplasts released from cells in culture by enzymatic digestion of cell walls to the elicitors.

In order to specifically gain access to the nuclear proteome, nuclei were isolated from the protoplast incubated at 18 °C for 16 hours under 3 conditions: 1 µM flg22, 1 µM nlp20 and untreated in case of control. This was repeated twice for a total of three independent experiments. The cellular suspension resulting as a product of the isolation procedure was also retained and used to prepare the cellular protein fraction comprising all proteins with the exception of those in the nucleus. The isolated nuclei were characterized by fluorescence microscopy after DAPI staining which attested to their successful isolation in an intact and round form (Figure 1A). Nuclear proteins were extracted and in-solution digested with trypsin along with the cellular proteins. The dissolved peptides were analyzed with liquid chromatography mass spectrometry (LC-MS) using a Data-Dependent Acquisition (DDA) scan strategy. The MS analysis identified 3899, 3212 and 3081 protein groups (set of proteins identified with a non-redundant peptide set, hence referred to as proteins) in the nuclear fraction of untreated, flg22 treated and nlp20 treated samples, respectively. 5633, 4742 and 5636 proteins were identified in the cellular fractions of the respective samples if they were identified in at least one of the replicate experiments (Supplementary file 1-Tables 1-3 and 7-9). The overlap between the fractions was 2587, 2301 and 2252 proteins respectively (Figure 1B).

**Figure 1.**
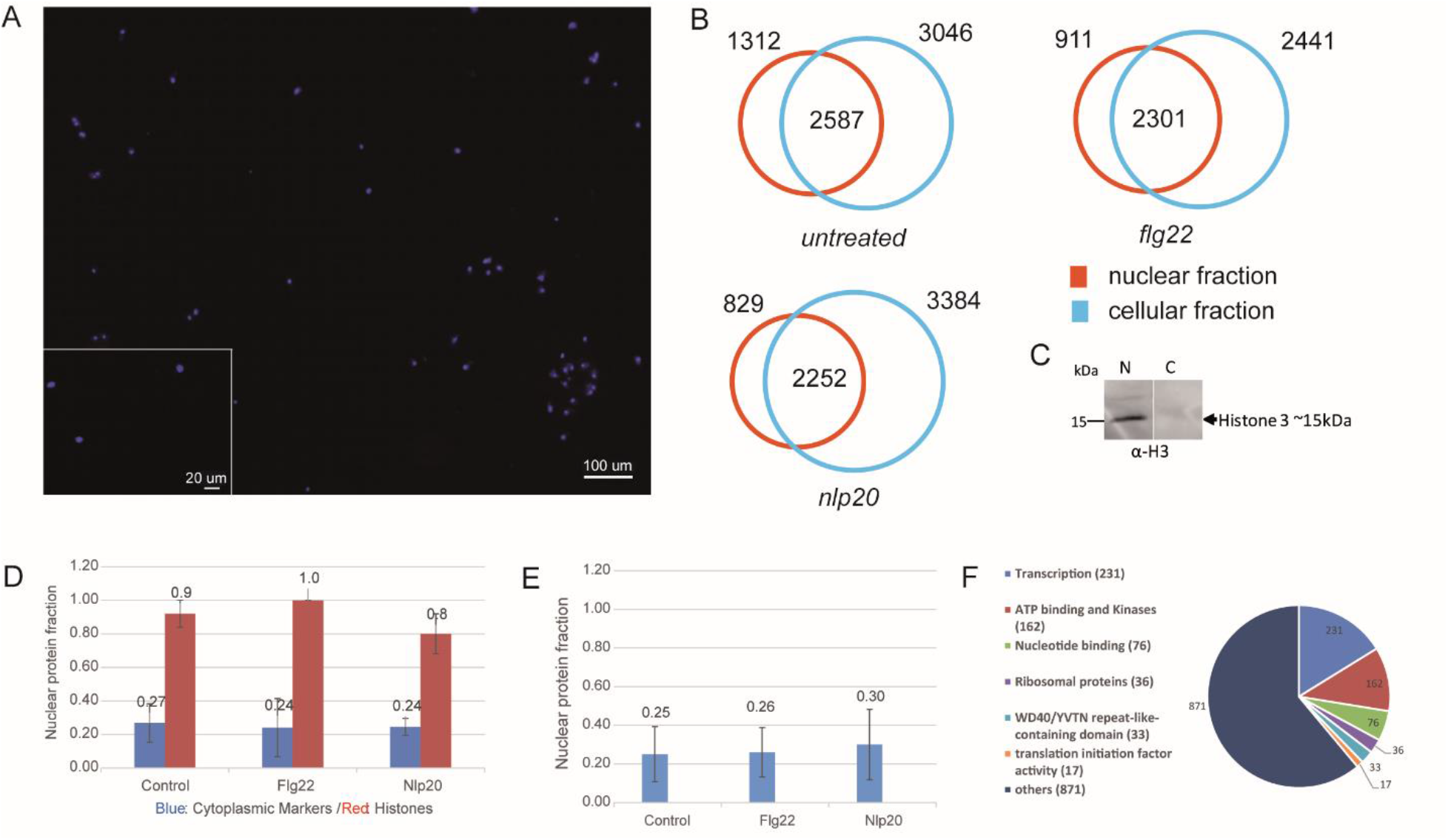
Extraction of the nuclear proteome. A. DAPI staining fluorescence microscopy of nuclei isolated from dark grown Arabidopsis thaliana cell culture. B. Total proteins identified in nuclear and cellular fractions in three independent experiments in untreated, flg22 and nlp20 treated cell cultures. C. Western blot of nuclear and cellular protein fraction with anti-Histone H3 antibody, respective Mw range is shown. D. Median nuclear enrichment scores (Npfn values) in three independent experiments of all identified histone proteins and of cytoplasmic markers (2 cytoplasmic markers were absent in shared proteins in flg22 and nlp20 conditions and were not used in both conditions). Error bars denote median absolute deviation. E. Median Npfn values of mitochondrial and Golgi markers. Error bars denote median absolute deviation. F. DAVID gene ontology classification of 1426 proteins of the untreated nuclear protein fraction annotated as nuclear proteins by DAVID bioinformatics tool.

### Defining the *Arabidopsis thaliana* nuclear proteome

A central issue in all organelle isolation procedures is the purity and integrity of the preparation. Conventionally the purity of the extracted nuclear proteome is assessed by way of western blot of nuclear protein markers, often histones. We also did this using antibody against histone H3 (Figure 1C) but in addition devised an approach to assess quality directly from the MS data. We calculated the fraction of each protein’s abundance in the nuclear and cellular protein fractions independently by way of the acquired MS data as mentioned in the methods section under collective data analysis. This can be interpreted as an enrichment score which in brief is the ratio of a protein’s MS signal in the nuclear or cellular fraction to its total MS signal in both fractions. Thus exclusive detection in the nuclear fraction would give a nuclear enrichment score (Npf_n_) of 1 whereas the score would be 0 if it were detected only in measurements of the cellular fraction. The score for the cellular fraction (Cpf_n_) would be the inverse.

The median nuclear enrichment score (Npf_n_) of all histones in the control, flg22 and nlp20 samples is shown in Figure 1D (for each histone see Supplementary file 1, Tables 4-6). All samples show enrichment of histones indicating successful isolation of nuclei and extraction of the nuclear proteome. In addition, Npfn values of Sad1/UNC84 homology (SUN2), KAKU4, WIP3, WiT1 and MAD1/NES1, five nuclear envelope / inner nuclear membrane proteins, were equal to 1 indicating exclusive presence in the nuclear protein fraction. To get an impression of the extent of inevitable contamination of the experimental nuclear proteome by cellular proteins, the fraction of the abundance of nine *bonafide* cytoplasmic markers (phosphoenolpyruvate carboxylase 1, phosphoenolpyruvate carboxylase 2, phosphoenolpyruvate carboxylase 3, Actin 1-3, Actin 7, Actin 8, Actin 12, sucrose phosphate synthase 1F and sucrose phosphate synthase 2F) were used. In contrast to histones, the median nuclear enrichment score (Npf_n_) of these markers was low (Figure 1D). In addition we calculated median Npf_n_ values for the known mitochondrial and golgi apparatus markers, voltage dependent anion channel 1, 2 and 3 as well as isocitrate dehydrogenase 1 and subunit 2 (mitochondrion) and coatomer gamma-2 subunit (golgi apparatus) which also were in the range of the cytoplasmic markers (Figure 1E). FD-GOGAT and FNR1 and 2 which are known plastid markers were completely absent from nuclear fractions. Together these results show that the isolation of nuclei was largely free of cytoplasmic contaminants and co-purifying organelles and debris and that the extracted nuclear proteome was of high purity.

Regarding the nuclear proteome, the proteins shared by both nuclear and cellular fractions (Figure 1B, intersections) may indeed be common to both and underlie some type of trafficking between nucleus and other organelles or cytoplasm or may simply be inevitable experimental contaminations of the nuclear fraction. To address this issue, we decided to use the median Npf_n_ values of the cytoplasmic markers (0.27, 0.24 and 0.24 respectively) described above as arbitrary cut off limits to define contamination in the three biological conditions and produce a curated set of nuclear proteins. All proteins with Npf_n_ values higher than this cut off limit were considered genuine nuclear proteins whereas those proteins with Npf_n_ values lower than the cut off limit were considered cellular protein contaminants and were discarded. The used cytoplasmic markers were discarded. This led to a curated set of nuclear proteins consisting of 2839, 2259 and 2096 proteins under control, flg22 and nlp20 conditions respectively that we define as the nuclear proteome under these biological conditions (Supplementary file 1-Tables 4-6).

To further validate the nuclear proteomes, we analyzed the curated protein lists with several bioinformatics tools that either predict protein sub-cellular localization or contain experimental evidence of it. These were the DAVID Bioinformatics resources 6.8 gene ontology tool (Huang et al., 2009a; Huang et al., 2009b), the *Arabidopsis* protein subcellular localization database (SUBA4) (Hooper et al., 2017), and LOCALIZER (Sperschneider et al., 2017), a software that predicts organelle subcellular localization by searching for targeting sequences such as NLS in protein primary structure. We also compared our experimentally determined proteomes with previously published nuclear and sub-nuclear proteomes (Bae et al., 2003; Bigeard et al., 2014; Calikowski et al., 2003; Chaki et al., 2015; Goto et al., 2019; Mair et al., 2019; Palm et al., 2016; Pendle et al., 2005; Sakamoto and Takagi, 2013). As a result 89 % of the nuclear proteins in each condition consisted of either experimentally verified nuclear proteins, proteins annotated to the nucleus or predicted to be localized in the nucleus (redundancy was removed), (Supplementary file 1-Tables 4-6). This underscores the high quality of our nuclear proteome preparation.

The DAVID bioinformatics tool was used to further annotate the nuclear proteins and classify them according to their function. 1426 proteins of the proteome measured under untreated conditions were classified initially as belonging to the nucleus. This set was then further input into DAVID. These proteins were categorized into 83 clusters and 6 main protein classes (Figure 1F and Supplementary file 1-Table 10). The 6 main classes were: transcription (231 protein), ATP binding & kinases (162 proteins), nucleotide binding (76 proteins), ribosomal proteins (36 proteins), WD40 (33 proteins) and translation initiation factor activity (17 proteins). Proteins annotated as related to the process of transcription, i.e. transcription factors and transcriptional co-activators were classified into families, as shown in Supplementary file 1-Table 11. The top 3 transcription factors families pertaining to the number of proteins identified were bZIP (13 proteins), WRKY (8 proteins) and Trihelix (8 proteins). Thus our LC-MS measurements of the nuclear protein fraction allowed insight into the transcription factor landscape in the nucleus which attests to the sequencing depth of our method. All together our results constitute a highly complete quality catalog of the nuclear proteome of *Arabidopsis thaliana* cell culture under homeostasis and induced immunity.

### Protein import into the nucleus under flg22 and nlp20 stimuli

We compared the curated nuclear protein sets identified under control, flg22 and nlp20 conditions as shown in Figure 2A. 1525 proteins were common to all the three conditions, but 269 and 223 were specific to flg22 and nlp20 respectively. This means that in our experiments considering our detection limits these proteins appeared in the nucleus only after elicitation of immunity with one of the two elicitors respectively and therefore they were likely imported. In order to investigate this further, we first checked the cellular proteins measured under the non-elicited control conditions for the presence of these specific proteins. 157 out of the 269 proteins appearing in the nucleus after flg22 exposure and 73 out of the 223 proteins appearing after nlp20 exposure were also measured in the cellular fraction without elicitation. Secondly, the nuclear enrichment (Npf_n_) and cellular enrichment scores (Cpf_n_) of the proteins were compared between control and flg22 and nlp20 elicited samples for the two sets of putatively imported proteins (157 and 73 proteins respectively). The Cpf_n_ decreased in both sets upon induction of immunity with either flg22 or nlp20 when compared to untreated samples. Strikingly, the Npf_n_ increased proportionally in elicited samples when compared to control (Figure 2B and C). The almost perfect proportionality and unitary sum is a strong indicator that these sets of proteins are indeed trafficked to the nucleus from the cytosol or some other cellular organelle upon elicitation of immunity wherein they play some particular function. Both sets of proteins are shown in Supplementary file 1-Table 12.

**Figure 2.**
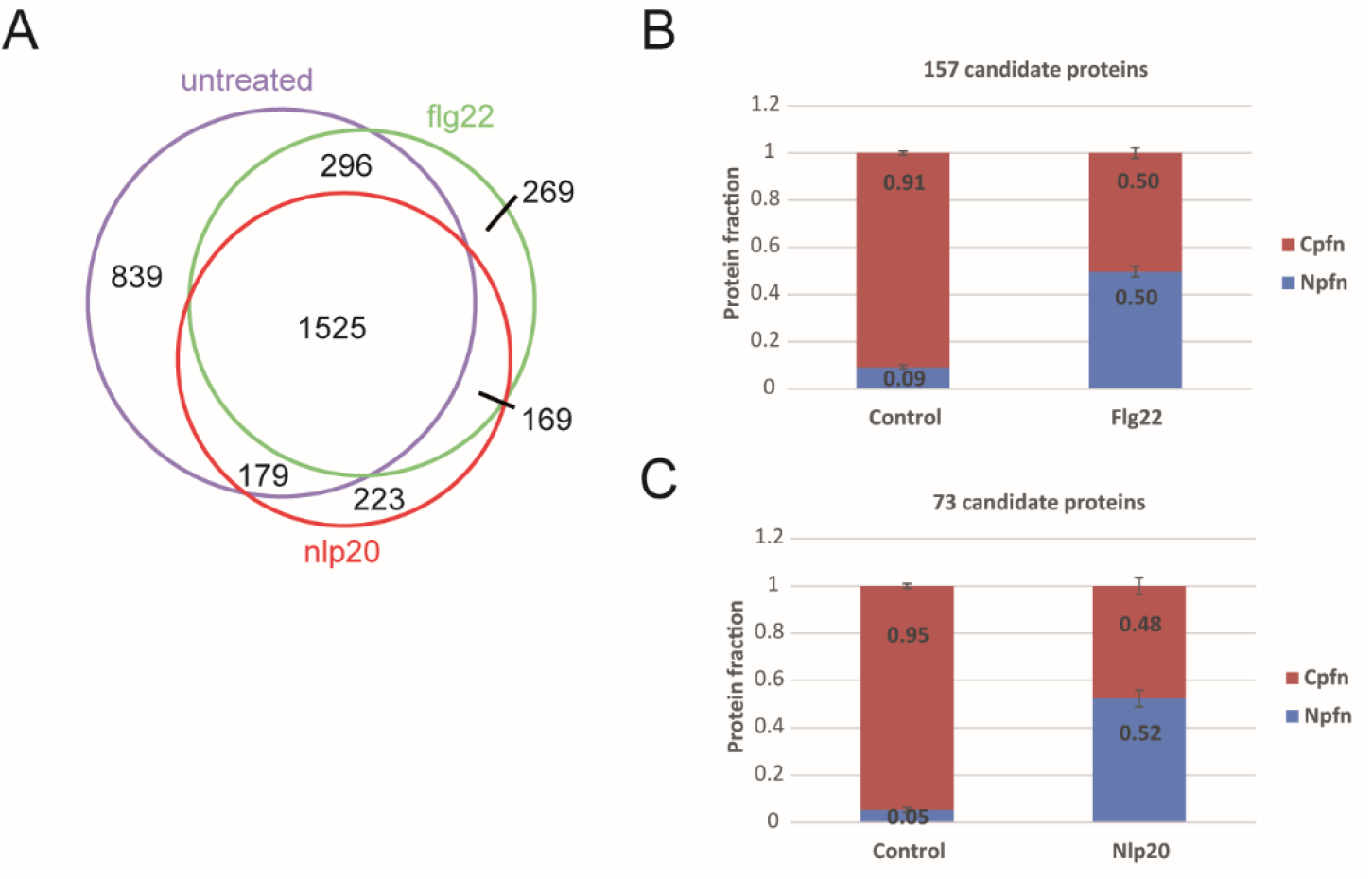
Protein import into the nucleus following elicitation of PTI. A. Intersections of curated nuclear protein fractions extracted from untreated and flg22 and nlp20 treated protoplasts. B. Mean nuclear and cellular protein enrichment scores (Npfn and Cpfn values) of 157 proteins identified in both the nuclear and cellular protein fractions without and following flg22 treatment. Error bars denote standard error. C. Mean nuclear and cellular protein enrichment scores (Npfn and Cpfn values) of 73 proteins identified in both nuclear and cellular protein fractions without and following nlp20 treatment. Error bars denote standard error.

### Comparison of nuclear proteomes under flg22 and nlp20 stimuli

To expand on the putative set of proteins imported into the nucleus following elicitation of immunity we were interested in identifying quantitative changes in protein abundance in the nuclear proteome in the three biological scenarios (control, flg22 and nlp20). To this end we looked at PSM count used as protein quantification index (PQI) in all three biological replicate experiments and performed multiple sample significance testing (ANOVA), with FDR multiples testing corrected significance threshold α=0.05, followed by post hoc test. 93 proteins showed significant change in their abundance between conditions (Supplementary file 1-Table 13). Hierarchical clustering of these proteins showed that the three biological scenarios produced specific clusters (Figure 3A). The protein dendrogram was divided into 4 main clusters as follows: proteins with decreased abundance in the nucleus following either flg22 or nlp20 stimulus (cluster 1), proteins with increased abundance in the nucleus following nlp20 stimulus (cluster 2), proteins with increased abundance in the nucleus following nlp20 and flg22 stimulus (cluster 3), proteins with increased abundance in the nucleus following flg22 (cluster 4) (Figure 3A and Supplementary file 1-Table 14). The 4 clusters comprise of proteins showing significant change in their abundance comparing elicited immunity to control. The proteins in these 4 clusters should play potential physiological roles in the nucleus during Pattern-triggered immunity (PTI). The 93 statistically significant proteins were annotated as being involved in several cellular processes such as mRNA processing, nucleotide binding, rRNA binding and include some protein families such as ribosomal proteins and prohibitin proteins.

**Figure 3.**
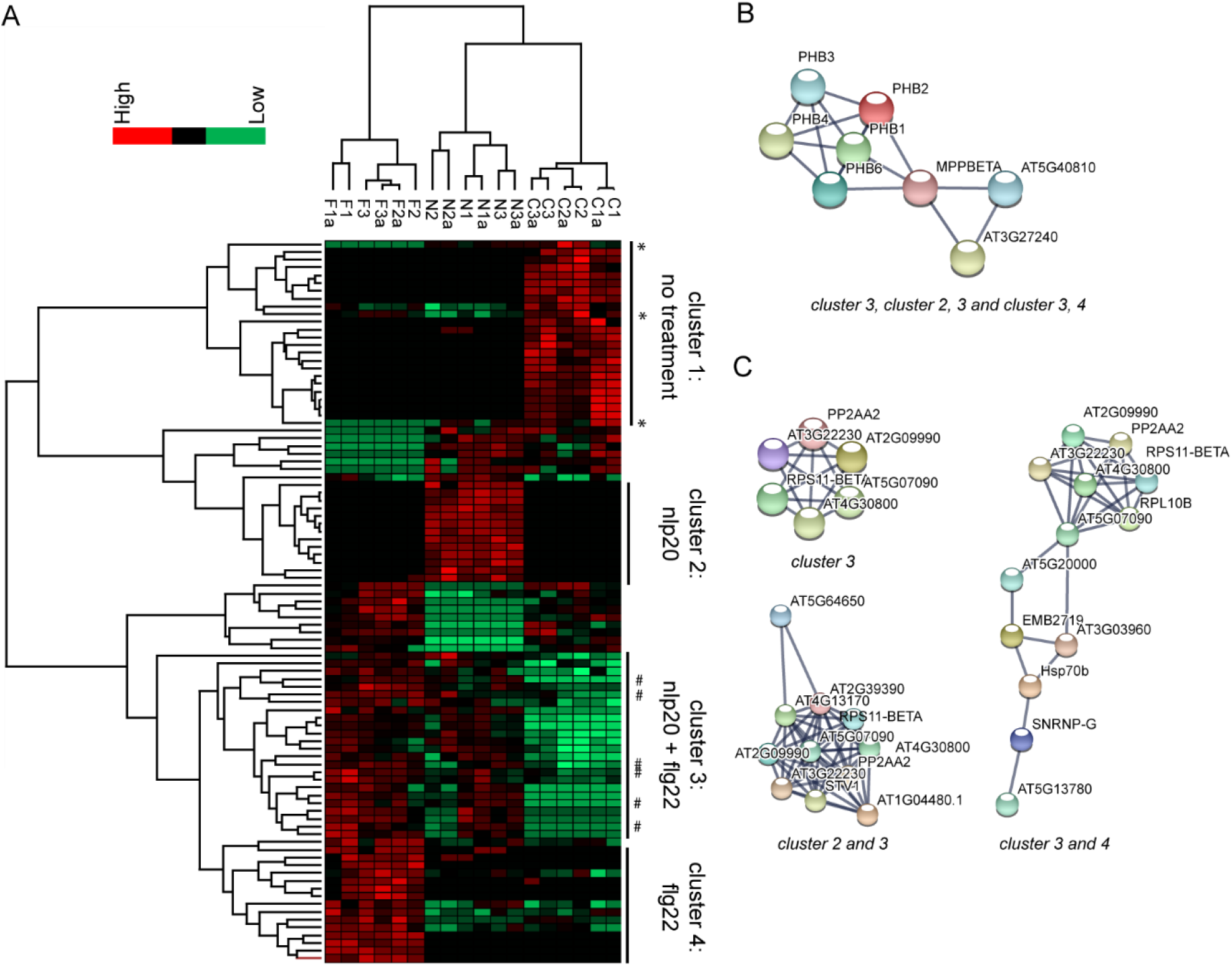
93 proteins showing significant changes in their abundance in the nuclear protein fraction following flg22 or nlp20 treatment. Multiple sample significance testing (ANOVA) was done, with FDR multiples testing corrected significance threshold α=0.05. A. Hierarchical cluster analysis (HCL) shows clustering of samples according to sample type. Rows represent proteins. PQI values were z-score transformed. Proteins are colored according to their abundance. * denotes decrease in abundance under effect of only flg22 or nlp20. # denotes increase in abundance under effect of flg22 only. B. STRING database binary protein interaction network of Prohibitins with Cytochrome C generated when indicated clusters were used as input sets. C. STRING database interaction networks of ribosomal and associated proteins when indicated clusters were used as input sets.

The proteins whose abundance increased upon stimulus with an elicitor (proteins in clusters 2, 3 and 4) were input into the STRING protein interaction database to identify potential physical interactions between them and putatively infer function of proteins in complexes. The abundance of all of the members of the prohibitin family which are prohibitins 1, 2, 3, 4 and 6 increased significantly upon treatment with both flg22 and nlp20 (Supplementary file 1-Table 14) and all interacted physically with one another. This prohibitin complex was expanded by three members of the mitochondrial bc_1_complex, MPPBETA (AT3G02090) and Cytochrome C1 family proteins AT5G40810 and AT3G27240 (Figure 3B). These proteins are all found in the mitochondrion and our results present evidence that they are translocated to the nucleus upon PTI induction.

19 ribosomal proteins showed a significant change in their abundance in between untreated and flg22 and nlp20 treated samples. 10 of these increased significantly in their abundance whereas the abundance of 9 decreased significantly (Supplementary file 1-Table 14). The abundance of 3 ribosomal proteins (S4, S5, L27e) was elevated upon either flg22 or nlp20 treatment. Conversely, the abundance of 5 ribosomal proteins (L13, L14p, L17, L24e, L29) increased specifically upon exposure of the protoplasts to nlp20 and the abundance of 2 ribosomal protein (L16p and S11) specifically upon exposure to flg22. Protein interaction analysis showed a core set of ribosomal proteins interacting in both PTI scenarios (Figure 3C top left panel). This core cluster however differentially expanded as proteins responding only to flg22 or nlp20 were added to the common input set (Figure 3C bottom left and right panels). This was particularly pronounced following elicitation of PTI by flg22 (Figure 3C right panel). It has been reported by us and others (see Bassal 2020 for citations) (Bassal et al., 2020) that ribosome composition is promiscuous dependent on cellular state and our result imply the same in the context of ribosome assembly in the nucleus. Finally, the abundance of another protein of interest to us, phospholipase D alpha 1 also increased significantly following stimulus of protoplasts with either flg22 or nlp20.

## Discussion

### Defining the nuclear proteome

In this study the *Arabidopsis thaliana* nuclear proteome was investigated following treatment of protoplasts cultured in the absence of light with two elicitors of PTI, flg22 and nlp20. Despite the central role of the nucleus in regulating gene expression, the plant nuclear proteome remains somewhat understudied (Narula et al., 2013; Yin and Komatsu, 2016). An early study using two-dimensional gel-electrophoresis (2-DE) characterized the nuclear proteome upon cold stress but the coverage was limited (Bae et al., 2003). Several studies have reported nuclear sub-proteomes such as nucleolar proteins, nuclear matrix or chromatin associated proteins and nuclear envelope (Bigeard et al., 2014; Calikowski et al., 2003; Chaki et al., 2015; Pendle et al., 2005; Sakamoto and Takagi, 2013; Tang et al., 2020). Also the nuclear proteome has been studied in tomato, potato, rice and barley (Howden et al., 2017; Narula et al., 2019; Petrovska et al., 2014; Rajamaki et al., 2020). In barley the authors employed fluorescence assisted cell sorting (FACS) to purify nuclei from plant tissue. Only recently however three studies have described the *Arabidopsis* nuclear proteome comprehensively, employing LC-MS (Goto et al., 2019; Mair et al., 2019; Palm et al., 2016). Mair et al. additionally used *in vivo* protein proximity biotin labeling to affinity purify nuclear proteins with high specificity. Our study complements these three, achieving a similar amount of proteins and thus similar comprehensive coverage in the context of plant immunity.

Organelle proteomes, including the proteome of the nucleus, are by definition smaller than the total cellular proteome which comprises around 10,000 proteins at any given time. Their qualitative composition, however may be substantially more dynamic because of biological context dependent protein in- and export. The use of the plant protoplast to isolate the nuclei was successful and produced high quality nuclei in a round and intact form (Figure 1A). Here we sought to gain a first impression of these molecular trafficking processes on a large scale by use of the LC-MS technology. Retrograde signaling from the chloroplast to the nucleus in plant immunity is well known. In our work we used cells cultured in the dark without chloroplasts to avoid the inevitable co-purification of these organelles with nuclei. Thus our study does not provide any insights into protein import from the chloroplast. The isolated nuclear fraction was of high purity and was enriched in histones and not in cytoplasmic or mitochondrial markers (Figure 1D and 1E). Indeed our results do shed light on protein import from these two compartments into the nucleus upon elicitation of PTI by both flg22 and nlp20.

An approach based on implementing a cut off limit was used to differentiate between nuclear and non-nuclear proteins. Generally cytoplasmic markers for instance actin and phosphoenolpyruvate carboxylase are abundant in the cytoplasm and not expected to be found in the nucleus (Dalmadi et al., 2019; Garcia et al., 2010; Genenncher et al., 2016). Therefore, the cytoplasmic markers can act as representatives for the non-nuclear protein contaminants when found in the nucleus and their nuclear protein fraction (Npf_n_) can be used as cut off limit to curate the nuclear proteins identified in the nuclear fraction. The purity of the curated nuclear proteins was verified and 89 % were either experimentally verified nuclear proteins, annotated to the nucleus or predicted to be localized in the nucleus. This reveals the success of the procedure and experimental approach used in identifying the nuclear proteome.

Proteins identified and annotated to the nucleus were involved in diverse nuclear functions. Transcription factors and transcriptional regulators control gene activity. Ribosomal proteins are part of the ribosomal biogenesis process (Turowski and Tollervey, 2015; Watkins and Bohnsack, 2012; Woolford and Baserga, 2013a). Nucleotide binding proteins (DNA or RNA binding), modulate gene expression and kinases are components of signal transduction cascades that regulate gene expression (Hucho and Buchner, 1997). Translation initiation factors act as a regulatory players in the translation process. WD40 proteins participate in various biological regulatory processes such as histone modifications, histone recognition and transcriptional regulation (Suganuma et al., 2008; Znaidi et al., 2004).

### Nuclear proteins import under flg22 and nlp20 stimuli

In this work the nuclear import was investigated under flg22 and nlp20 as triggers for immune response. Two lists of proteins were identified as potential candidates for import under both treatment conditions (157 proteins for flg22 condition and 73 proteins for nlp20 condition). Generally, protein transport into the nucleus is controlled by different mechanisms. Proteins smaller than 40-60 kDa are diffused in a passive manner but larger proteins need to be recognized by the nuclear transport receptors which bind the nuclear localization signals (NLS) on those proteins and facilitate import (Chook and Suel, 2011; Christie et al., 2016; Grossman et al., 2012; Mohr et al., 2009; Tamura and Hara-Nishimura, 2014; Timney et al., 2016; Wang and Brattain, 2007; Weis, 2003). In addition alternative mechanisms were also investigated (Guinez et al., 2005; Imamoto and Kose, 2012). In the flg22 list 33% of the proteins were predicted to have NLS, the molecular weight of the rest of the proteins in flg22 list were checked and 71% of them were less than 40-60 kDa. Similarly, in the nlp20 list 33% of the proteins were predicted to have NLS and 88% of the rest of the proteins were less than 40-60 kDa. Therefore, the proteins we postulate to be imported fulfill the requirements for the transport into the nucleus.

Proteins undergo their functions in a specific manner regarding time and location. Therefore, the designated proteins needed to be activated and recruited to certain subcellular locations only when required (Michaelson et al., 2008; Wiatrowski and Carlson, 2003). The activation could be through a signal transduction cascade that activates the protein by post translational modifications (Di Ventura and Kuhlman, 2016). In addition, nucleoporins regulate selectively the passage of certain stress-sensible proteins (Yang et al., 2017) by undergoing conformational changes upon immunoreceptor activation and allowing transport of specific macromolecules (Gu et al., 2016). In this work the nuclear import was investigated under flg22 and nlp20 stimulus, implying that the flg22 and nlp20 induced responses could directly or indirectly regulate nuclear import of the selected two sets of proteins by one of the above mentioned mechanisms.

### Comparison of nuclear proteomes under flg22 and nlp20 stimuli

The nuclear proteomes were investigated for quantitative changes in protein abundance in the three biological scenarios: untreated control, flg22 and nlp20 treatment. 93 proteins showed statistically significant changes in their abundance upon elicitation of one or both forms of PTI when compared to control. We will focus in the discussion on 2 proteins families and 1 protein and their potential roles in immunity: ribosomal proteins, prohibitin proteins and phospholipase D alpha 1.

### Ribosomal proteins

The ribosomes are the cellular machines required for the process of protein synthesis. The maturation of the ribosomes is required for its function, the process of ribosome maturation is called ribosome biogenesis. Ribosome biogenesis involves association of ribosomal proteins with rRNA to constitute the ribosomal subunits (Sáez-Vásquez and Delseny, 2019; Thomson et al., 2013). Primary steps of ribosome biogenesis exist in the nucleus before exportation to the cytoplasm (Brown and Shaw, 1998; Henras et al., 2015; StępiĔski, 2014; Woolford and Baserga, 2013b). Adding to the function of ribosomal proteins in ribosome biogenesis and protein synthesis, they have various extra-ribosomal functions (Zhou et al., 2015) for example, transcription regulation and histone binding in the nucleus (Denmat et al., 1994; Dieci et al., 2009; Ni et al., 2006; Tchórzewski et al., 1999; Tu et al., 2011). Therefore, ribosomal proteins have been identified in the nucleus in many studies for instance in *Arabidopsis thaliana* (Chaki et al., 2015; Palm et al., 2016; Pendle et al., 2005). In this work ribosomal proteins were also identified in our nuclear proteome, 19 of them had a significant change in their abundance between control and immunity. Interestingly, these ribosomal proteins showed different abundance in the nucleus when treated with the two elicitors flg22 and nlp20 as mentioned in the results. These elicitor specific changes in abundance suggest that ribosomal proteins do not act similarly and have different functions in the two types of PTI. Accordingly, we can speculate that the ribosomal proteins with increased abundance in the nucleus under flg22 and nlp20 compared to control (10 proteins: 5 proteins specific to nlp20, 2 protein specific to flg22 and 3 proteins for both flg22/nlp20) play an active role in the nucleus during the immune response under different stimuli in *Arabidopsis*. On the other hand, the ribosomal proteins with decreased abundance in the nucleus under flg22 and nlp20 compared to control (9 proteins) have a repressed function in the nucleus during the *Arabidopsis* immune response. In a previous study the ribosomal protein transcripts were investigated in *Vanilla planifolia* when infected with *Fusarium oxysporum* (Solano de la Cruz et al., 2019). 7 ribosomal protein families that showed an increase in abundance in the nucleus after elicited immunity in our study also showed an increase in their transcript expression patterns in *Vanilla* after 2 days of *Fusarium* infection. These 7 protein families are: ribosomal protein L14p/L23e family protein, ribosomal L29 family protein, ribosomal protein L13 family protein, ribosomal protein L24e family protein, ribosomal protein L17 family protein, ribosomal protein S4 (RPS4A) family protein and ribosomal protein S5 domain 2-like superfamily protein. In addition, ribosomal protein L24e family protein was also detected exclusively in the nucleus of the *cerk1* background in *Arabidopsis* after chitosan treatment (triggering a MAMP-like response). The authors also observed that the ribosomal proteins were overrepresented after chitosan treatment (Fakih et al., 2016). This suggest that these 7 ribosomal protein families have distinct functions in plant immunity in different plants elicited by different pathogens and promiscuity of ribosomal proteins in ribosome assembly is known. This functional promiscuity is reflected by the different protein interactions undergone by the ribosomal proteins in the two PTI scenarios (Figure 3C).

### Prohibitins

Prohibitins are group of conserved proteins in eukaryotes including plants (Van Aken et al., 2010). They were reported to have several functions as scaffold proteins in apoptosis, mitochondrial biogenesis and the immune response (Yu, 2019). Prohibitins participate in plant defense response and in protection against stress, for example: they are involved in the rice defense response against fungus (Takahashi et al., 1999; Takahashi et al., 2003). PHB1 and PHB2 are localized in the mitochondria and participate in its biogenesis and in the plant response to stress in *Nicotiana benthamiana* (Ahn et al., 2006) and PHB3 is additionally localized to the chloroplast where it regulate the production of salicylic acid under UV and biotic stress in *Arabidopsis* (Seguel et al., 2018). Besides their localization in the mitochondria and chloroplasts, prohibitins have also been reported in the nucleus and act as transcription regulators in eukaryotes (Huang et al., 2019; Mishra et al., 2006; Peng et al., 2015; Thuaud et al., 2013). In *Arabidopsis thaliana* five prohibitins are expressed (PHB1, PHB2, PHB3, PHB4 and PHB6) (Van Aken et al., 2007) and all of them were identified in our nuclear proteome with increased abundance following treatment with both flg22 and nlp20. This indicates that the prohibitin family plays a crucial role in the *Arabidopsis* defense response in the nucleus. In previous studies, PHB2 was detected exclusively in the nucleus of *cerk1 Arabidopsis* plant after chitosan treatment (triggering a MAMP-like response) (Fakih et al., 2016) and prohibitin protein was also identified in the nucleus of *Solanum Lycopersicum* with increased abundance after 24 hour infection with *Phytophthora capsici* compared to non-infected plants (Howden et al., 2017). The results of these two studies support our findings of a probable role of prohibitins in the nucleus during the plant immune response. In addition, all five prohibitins interacted with each other and Cytochrome C (Figure 3B), another mitochondrial protein whose abundance also increased in the nucleus in PTI. Cytochrome C has been shown to have functions in the nucleus such as DNA damage repair and interaction with histone proteins (Gonzalez-Arzola et al., 2019) in addition to its well-known function in the mitochondrial respiratory chain. It is therefore tempting to speculate, that the prohibitins may act as a scaffold to traffic Cytochrome C from the mitochondrion to the nucleus in PTI.

### Phospholipase D alpha 1

We were able to identify the phospholipase D alpha 1 (PLD) protein in our nuclear proteome with increased abundance after treatment with both flg22 and nlp20 when compared to control indicating a possible role in the immune response within the nucleus. Generally, PLD1 is involved in many plant cellular process including the plant response to wounding, salt stress and plant-microbial interactions (Andersson et al., 2006; Bargmann and Munnik, 2006; Hong et al., 2010; Wang, 2005; Young et al., 1996; Zhao et al., 2013). Additionally, an early study reported that PLD1 was localized to the nucleus in mammalian cell lines and had a possible function in mRNA processing (Jang and Min do, 2011). Similarly in *Arabidopsis* PLD alpha and PLD gamma were also localized to the nucleus (Fan et al., 1999). Therefore, our findings are supported by previous studies which link PLD to the plant defense response against microbes as well as its nuclear localization. Recently, a research group reported the interaction between PLD alpha 1 and MPK3 (mitogen-activated protein kinase 3) in regulating salt stress in *Arabidopsis* (Vadovic et al., 2019). Mitogen-activated protein kinases play a significant role in plant defense response to biotic stress (Colcombet and Hirt, 2008; CristinaRodriguez et al., 2010; Latrasse et al., 2017; Šamajová et al., 2013) including MPK3 (Galletti et al., 2011) which was also localized to the nucleus (Vadovic et al., 2019). Hence, we can speculate on the possibility of a similar regulatory role of PLD alpha1 within the nucleus in the plant response to biotic stress upon interaction with MAPK or MPK3. Interestingly, Phospholipase D gamma 3 was identified in our nuclear proteome with decreased abundance after flg22/nlp20 treatment when compared to control, implying that different PLDs could have differing roles in nuclear associated processes in plant immunity.

## Acknowledgments

We thank the Leibniz Association for support and funding. Mohamed Ayash is funded by DFG 4006811449/GRK2498. Mohammad Abukhalaf is funded by DFG grant HO 5063/2-1. This work was also supported by de.NBI (FKZ 031 A 534A), a project of the BMBF (Bundesministerium für Bildung und Forschung). We thank Alexandra Gurowietz for support.

## Author contributions

W.H. conceived and oversaw the study. M. Ayash, M. Abukhalaf and W.H. designed experiments. M. Ayash, M. Abukhalaf, M.H., C.P. and D.T. performed experiments. M. Ayash, M.H., M.S. and W.H. analyzed data. M. Ayash and W.H. wrote the manuscript.

---

**Supplementary Figure 1.**
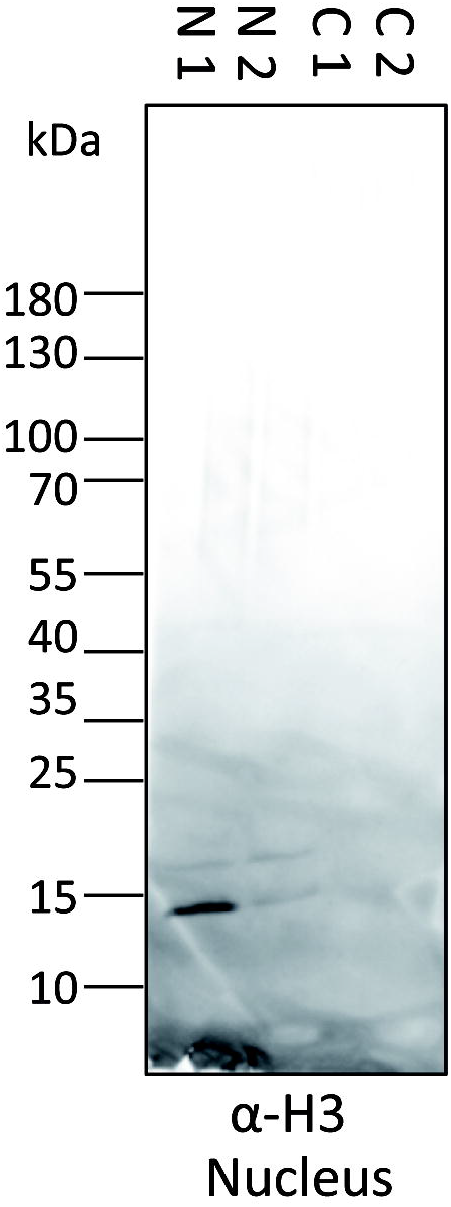
Western blot of nuclear and cellular protein fraction with anti-Histone H3 antibody. Two independent experiments are shown, N denotes nuclear and C cellular protein fractions.

